# Heterogeneity of synaptic NMDA receptor responses within individual lamina I pain processing neurons across sex in rats and humans

**DOI:** 10.1101/2023.08.09.550677

**Authors:** Annemarie Dedek, Emine Topcu, Christopher Dedek, Jeff S. McDermott, Jeffrey L. Krajewski, Eve C. Tsai, Michael E. Hildebrand

**Affiliations:** Department of Neuroscience, Carleton University, Ontario, Canada; Neuroscience Program, Ottawa Hospital Research Institute, Ontario, Canada; Lilly Research Laboratories, Indianapolis, Indiana, United States; Brain and Mind Research Institute, University of Ottawa, Ontario, Canada; Division of Neurosurgery, Department of Surgery, The Ottawa Hospital, Ontario, Canada

## Abstract

Excitatory glutamatergic NMDA receptors (NMDARs) are key regulators of spinal pain processing, and yet the biophysical properties of NMDARs in dorsal horn nociceptive neurons remain poorly understood. Despite the clinical implications, it is unknown whether the molecular and functional properties of NMDAR synaptic responses are conserved between males and females as well as from rodents to humans. To address these translational gaps, we systematically compared individual and averaged excitatory synaptic responses from lamina I pain-processing neurons of adult Sprague Dawley rats and human organ donors, including both sexes. By combining patch-clamp recordings of outward miniature excitatory postsynaptic currents with non-biased data analyses, we uncovered a wide range of decay constants of excitatory synaptic events within individual lamina I neurons. Decay constants of quantal synaptic responses were distributed in a continuum from 1-20 ms to greater than 1000 ms, suggesting that individual lamina I neurons contain AMPA receptor (AMPAR)-only as well as GluN2A-, GluN2B-, and GluN2D-NMDAR-dominated synaptic events. This intraneuronal heterogeneity in AMPAR– and NMDAR-mediated decay kinetics was observed across sex and species. However, we discovered a decreased relative contribution of GluN2B-dominated NMDAR responses as well as larger amplitude GluN2D-like events at human lamina I synapses compared to rodent synapses, suggesting species differences relevant to NMDAR subunit-targeting therapeutic approaches. The conserved heterogeneity in decay rates of excitatory synaptic events within individual lamina I pain-processing neurons may enable synapse-specific forms of plasticity and sensory integration within dorsal horn nociceptive networks.

## Introduction

Pain serves a critical role in alerting one to danger (Raja et al., 2020); however, despite this essential role, pathological forms of pain have resulted in an out-of-control clinical crisis. After decades of research and development, few substantial changes have been made to how pain is treated. Two major barriers that have obstructed a better understanding and treatment of pain are an overreliance on studies performed on male-only or unsexed animals (Mogil, 2012; Shansky & Murphy, 2021), and the translational species divide between rodent pain models and the target human clinical population (Gereau et al., 2014; Renthal et al., 2021).

The superficial dorsal horn of the spinal cord is a critical hub for nociceptive processing. It is the main site of entry for high-threshold nociceptive primary afferents (Peirs & Seal, 2016; Todd, 2010), as well as the site of processing and integration by local interneuron networks (Chamessian et al., 2018; Gutierrez-Mecinas et al., 2016; Todd, 2017) that also receive descending efferent input from the brain (Ossipov et al., 2014). Lamina I, the outermost region of the dorsal horn, is comprised of interneurons as well as ascending projection neurons and is thus a key determinant of the output of spinal nociceptive circuits (Choi et al., 2020; Hachisuka et al., 2020; Häring et al., 2018).

With an increase in sex-inclusive preclinical pain research (Mogil, 2020), striking differences in neuroimmune signalling and downstream mediators of spinal hyperexcitability have been found between male and female rodent models of chronic pain (Chen et al., 2018; Mapplebeck et al., 2018; Sorge et al., 2015). We have recently discovered a sexually dimorphic mechanism that drives the potentiation of excitatory NMDA receptors (NMDARs) at lamina I synapses in both rat and human tissue models of pathological pain (Dedek et al., 2022).

However, it remains unclear to what degree the molecular and functional properties of NMDAR signalling are conserved between sexes in baseline physiological conditions, particularly in the human spinal cord.

Recent translational studies have increased our understanding of the genetic underpinnings of human nociception by creating atlases of single cell gene expression for both human dorsal root ganglia (DRG) (Tavares-Ferreira et al., 2022) and spinal cord (Yadav et al., 2023) tissue. Yet, studying the functional properties of intact human nociceptive neurons with electrophysiology has proven difficult. Surgical excision of brain regions in treatment-resistant cases of epilepsy has enabled researchers to gain insight into the electrophysiological properties of neurons in the human brain (Howard et al., 2022; Kushner et al., 2022; Malkin et al., 2022), but these samples are limited to pathological cases and do not provide insight into potential species differences in resting-state physiology. In the case of the spinal cord, no such surgical excision models exist, but our recently developed *ex vivo* assays using human organ donor tissue samples have opened the door for the electrophysiological characterization of dorsal horn nociceptive neurons (Dedek et al., 2019).

Excitatory NMDARs are central determinants of synaptic plasticity. Four different genetically encoded GluN2 subunits (GluN2A-D) create heterogeneity in both the biophysics and regulation of glutamate-activated NMDAR currents (Paoletti et al., 2013). For example, GluN2A-, GluN2B/C-, and GluN2D-containing receptors mediate NMDAR currents with relatively fast, moderate, and slow deactivation rates, respectively, enabling different degrees of temporal summation and integration. Within individual neurons of male or unsexed rodent brain slices, distinct GluN2 subunits have been proposed to drive synapse-specific NMDAR biophysical properties and associated mechanisms of plasticity (Fleidervish et al., 1998; Ito et al., 2000; Kumar & Huguenard, 2003; Otmakhova et al., 2002). However, this intraneuronal heterogeneity in synaptic NMDARs remains to be investigated across sex and species and has not been considered for neurons in the highly interconnected dorsal horn nociceptive network.

Here, we combined an unbiased analysis approach with electrophysiological recordings of AMPAR– and NMDAR-mediated miniature excitatory postsynaptic currents (mEPSCs) to characterize the functional contributions of these receptors’ subunits to excitability at individual lamina I synapses. To address the translational gap between preclinical rodent models and humans, we included patch-clamp recordings of lamina I neurons from viable slices of both rat and human organ donor spinal tissue. We validated our biophysical analysis approach with pharmacological experiments using GluN2 subunit-specific antagonists. In combination, this experimental strategy provides a comprehensive and direct comparison of excitatory synaptic responses for individual spinal cord pain signalling neurons across sex in both rats and humans.

## Results

### Patch-clamp recordings from male and female rat and human lamina I neurons reveals heterogeneity between individual excitatory synaptic responses

Although NMDARs play well-documented roles in modulating excitability in spinal nociceptive circuits (Hildebrand et al., 2016; Xie et al., 2016; Yamamoto & Yaksh, 1992; Zhou et al., 2019), our knowledge of NMDAR biophysical properties within dorsal horn neurons is based almost entirely on studies performed on male or unsexed rodents (Dedek & Hildebrand, 2022). This is particularly important since we recently discovered divergent mechanisms of dorsal horn NMDAR dysregulation by sex in both rats and humans (Dedek et al., 2022). We therefore systematically compared baseline synaptic NMDAR activity across sex and species by recording outward mEPSCs in lamina I neurons from *ex vivo* adult rat and adult human spinal cord slices of both sexes (Figure 1A). Following two minutes of acclimatization, we recorded pharmacologically isolated mEPSCs at +60 mV (Hildebrand et al., 2014) for a standardized 8-minute time interval. We then detected and aligned all non-overlapping mEPSCs to analyze the peak amplitudes and decay constants (τ_decay_) of these excitatory synaptic events (Figure 1A).

**Figure 1.**
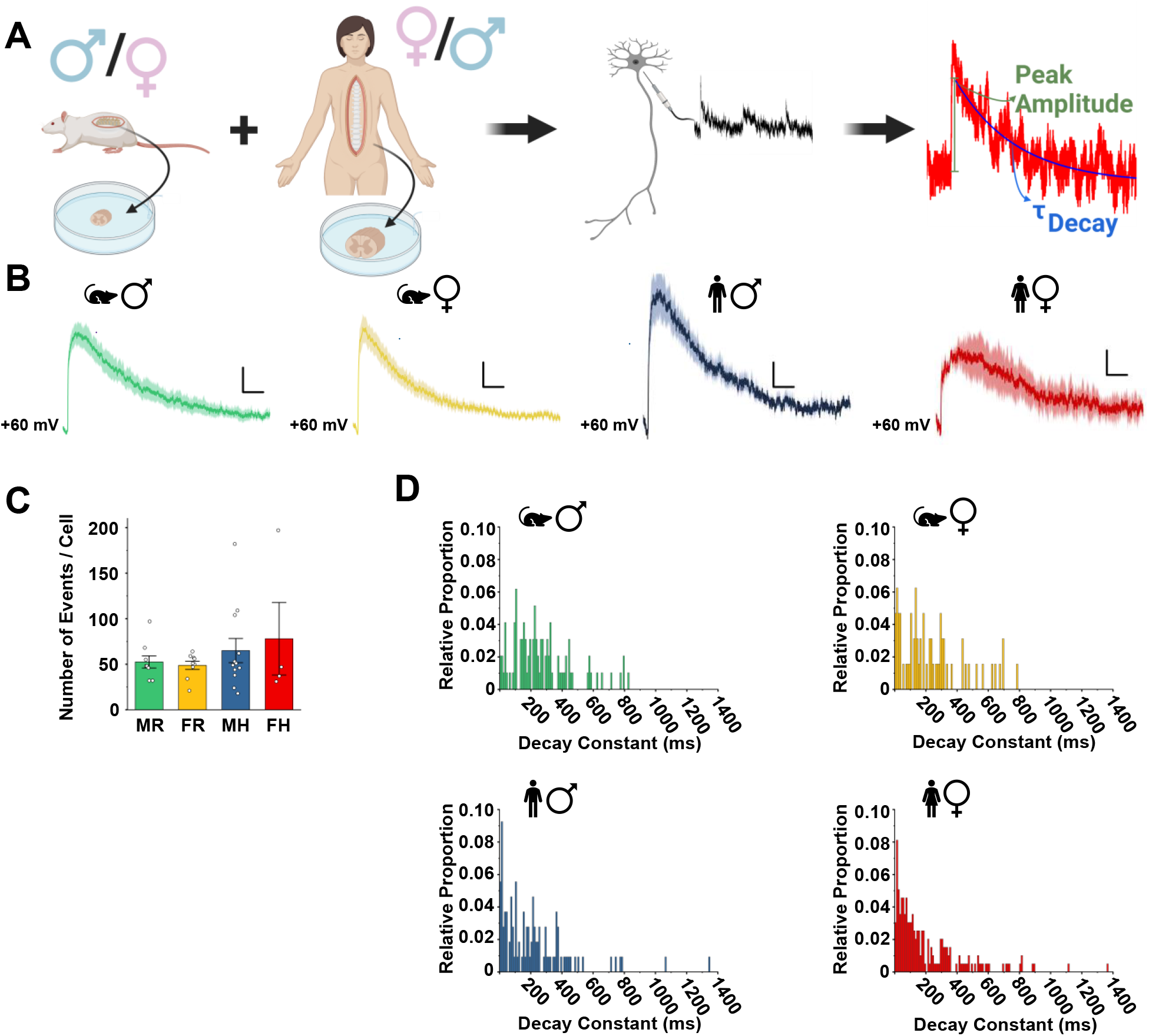
There is substantial variability between individual mEPSC events within lamina I neurons of male and female rats and humans. **A**) Experimental approaches in this study. *Ex* vivo spinal cord slices were prepared from male and female adult rats and humans. After establishing the whole cell patch clamp configuration on lamina I pain processing neurons, mEPSCs were recorded at +60mV. Individual mEPSC events were analyzed to measure the peak amplitude and decay constant (τ_decay_). Created using BioRender. **B**) Averaged mEPSCs for male and female rats and humans using previous GluN2B-focussed analysis criteria (Dedek et al., 2019) appear to have similar properties. Male rat n = 9, female rat n = 9, male human n = 7, female human n = 2. Scale bars = 100 ms (x-axes); 5 pA (y-axes) **C**) Using a new, unbiased criteria, a sufficient number of events was detected for all analyzed neurons, and the number of mEPSCs detected during an 8-minute recording period for each cell was not significantly different across all groups. Male rat n = 9, female rat n = 9, male human n = 12, female human n = 4. **D**) As shown here for a representative neuron in each group, there was a wide distribution of mEPSC decay constants within individual male (green, top left) and female (yellow, top right) rat as well as male (blue, bottom left) and female (red, bottom right) human lamina I neurons.

In our first characterization of NMDAR responses at lamina I synapses, we identified that GluN2B-containing receptors dominate NMDAR mEPSCs in adult male rats (Hildebrand et al., 2014). Thus, we subsequently used GluN2B-focussed mEPSC analysis techniques to study the regulation of GluN2B-mediated synaptic NMDAR responses across species and sex (Dedek et al., 2019, 2022). Using the GluN2B-focussed analysis criteria here, which include a constrained range of mEPSC decay rates and peak amplitudes (see Methods), we generated averaged mEPSC traces for all treatment groups (Figure 1B). Visual inspection of these lamina I neuron mEPSCs appeared to indicate that synaptic NMDAR properties are relatively conserved from rats to humans for both males and females, with a potential decreased amplitude for female human neurons (Figure 1B). However, this restrictive analysis criteria meant that many mEPSC events were excluded as they decayed too fast or too slow or had amplitudes that were below the 10 pA minimum or above the 200 pA maximum. Interestingly, in contrast to male and female rats where all recorded cells could be included (n = 9 cells each), only 7 of 12 neurons from male humans and 2 of 4 neurons from female humans had a sufficient number (>20) of acceptable events to be included in these averaged plots using the original analysis criteria (Figure 2B). To address this, we developed new unbiased analysis criteria that encompasses all potential AMPAR– and NMDAR-mediated mEPSCs that exhibit decay within 4000 ms and that are of sufficient signal-to-noise threshold (see Methods). Using these new criteria, a sufficient number of acceptable mEPSC events was identified for analysis for all cells in all four sex/species groups (Figure 1C). Although there was considerable variation from cell-to-cell, the number of analyzed mEPSC events was always above 18 and did not significantly differ across sex or species (53 ± 7 events in 8 male rats, n = 9 neurons; 49 ± 4 events in 8 female rats, n = 9 neurons; 65 ± 13 events in 5 male humans, n = 12 neurons; and 78 ± 40 events in 3 female humans, n = 4 neurons; Figure 1C).

**Figure 2.**
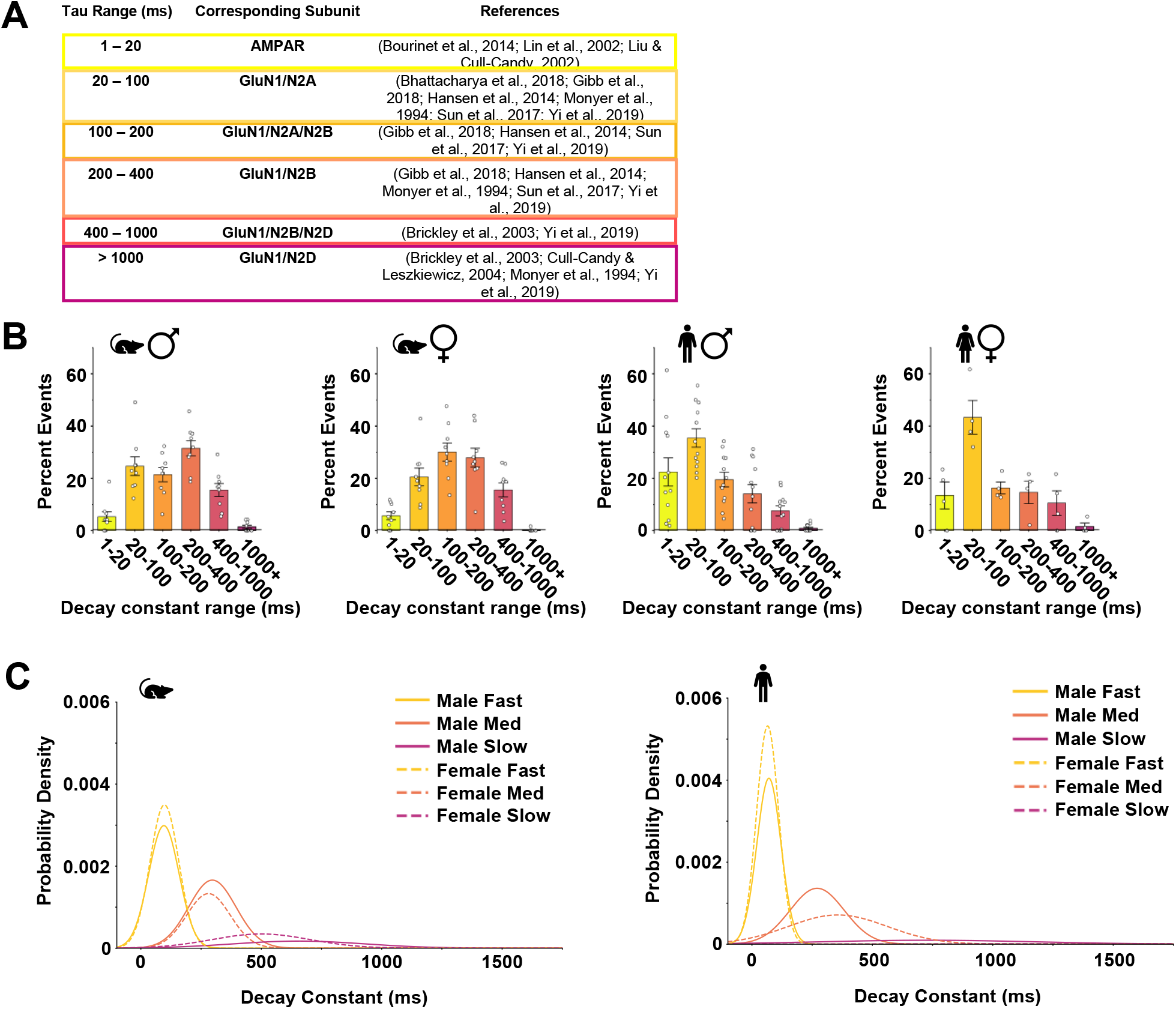
Intraneuronal heterogeneity of mEPSC decay constants is conserved across sex and species, with a decreased proportion of GluN2B-like mEPSCs in humans compared to rats. **A**) Measured decay constants (τ_decay_) for each mEPSC were binned according to tau-decay ranges that correspond to specific AMPAR– and GluN2-NMDAR subunit-mediated responses, as reported in the cited literature. **B)** Percent of mEPSC events in each decay constant bin for recordings on lamina I neurons from male rats (left, n = 9), female rats (middle left, n = 9), male humans (middle right, n = 12) and female humans (right, n = 4). A two-way ART ANOVA within species revealed no statistically significant differences between sex, however, the percent frequency of events was not distributed equally across decay constant range (see Tables 1 and 2 for lists of pairwise comparisons). **C)** Gaussian distributions outlining the probability density of all mEPSC decay constant values for all cells in male (sold lines) and female (dashed lines) rats (left) and humans (right).

**Table 1.**
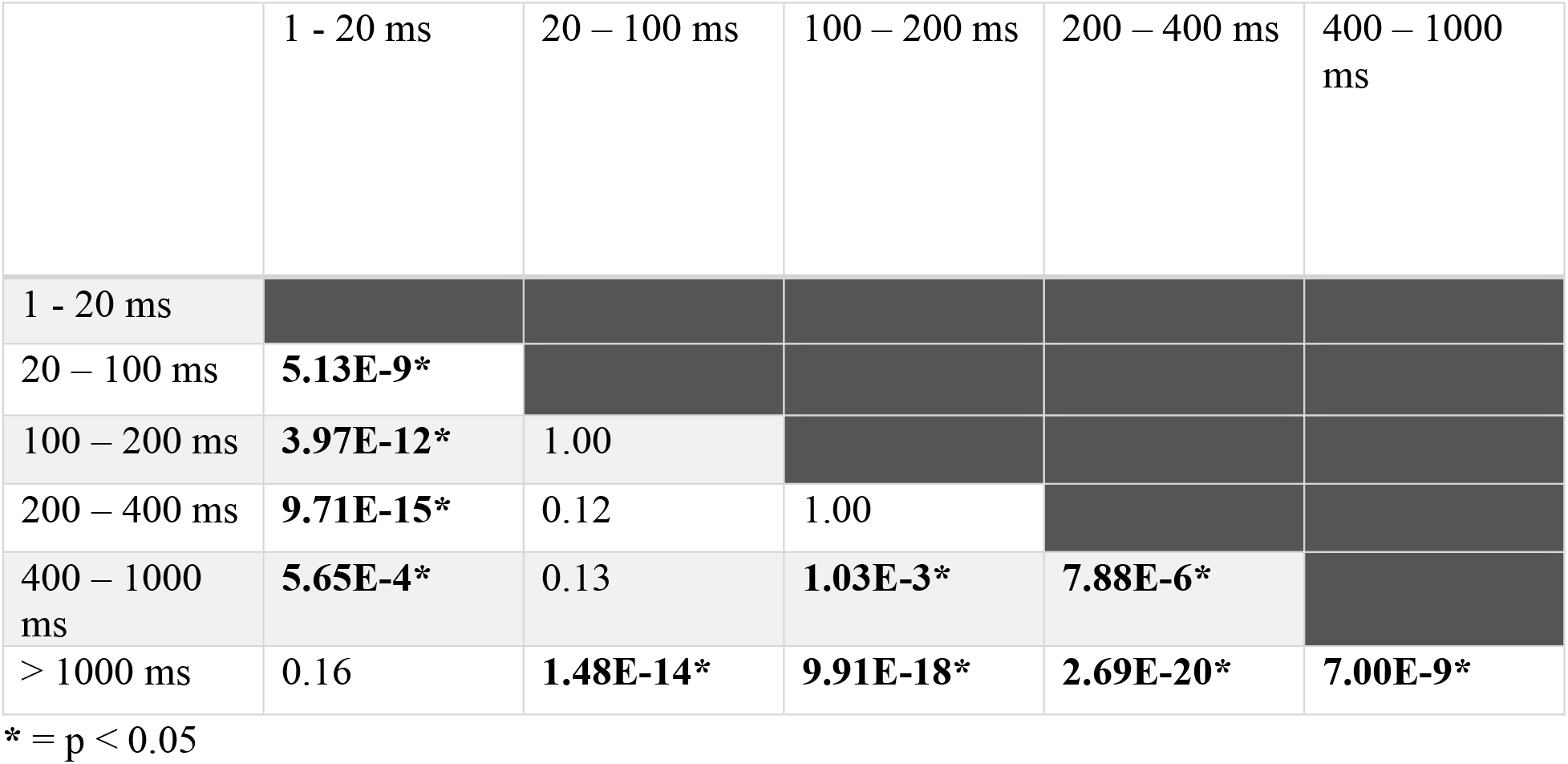
Statistical significance of within-sex Bonferroni pairwise comparison between the percent frequency of mEPSC events in each of the τ_decay_-defined bins in rat lamina I neurons.

**Table 2.** Statistical significance of within-sex Bonferroni pairwise comparison between the percent frequency of mEPSC events in each of the τ_decay_-defined bins in human lamina 1 neurons.

|  | 1 - 20 ms | 20 – 100 ms | 100 – 200 ms | 200 – 400 ms | 400 – 1000 ms |
| --- | --- | --- | --- | --- | --- |
| 1 - 20 ms |  |  |  |  |  |
| 20 – 100 ms | **2.53E-3\*** |  |  |  |  |
| 100 – 200 ms | 1.00 | **5.15E-3\*** |  |  |  |
| 200 – 400 ms | 1.00 | **1.50E-5\*** | 1.00 |  |  |
| 400 – 1000 ms | 0.083 | **2.27E-8\*** | **0.045\*** | 1.00 |  |
| > 1000 ms | **4.00E-6\*** | **6.50E-14\*** | **1.00E-6\*** | **7.46E-4\*** | 0.10 |

\* = $p < 0.05$

Upon closer visual inspection of individual mEPSCs identified through the unbiased analysis approach, we observed a large degree of heterogeneity between mEPSCs within single lamina I neurons. This heterogeneity between unitary mEPSC events is masked by the averaged mEPSC plotting approach shown in Figure 1B and by neuronal population-based analyses used in our past publications (Dedek et al., 2019, 2022; Hildebrand et al., 2014, 2016). To quantify this variability, we fit the decay component of individual mEPSC events and plotted the distribution of mEPSC decay constants. As shown for representative neurons from each of the four groups in Figure 1D, we found a wide distribution of mEPSC decay constants within individual lamina I neurons from male and female rats and humans.

### A range of decay constants, corresponding to distinct AMPAR– and NMDAR subunit-dominated synaptic responses, is conserved between male and female lamina I neurons for both rats and humans

Given the observed heterogeneity in decay rates of discrete mEPSC events within individual lamina I neurons, we set out to systematically compare the relative distributions of mEPSC decay constants by cell for both sexes and species. We grouped these mEPSC decay constants into discrete bins that corresponded to previously published ranges of decay constants for AMPAR-mediated, GluN1/GluN2A-mediated, GluN1/GluN2A/GluN2B-mediated, GluN1/GluN2B-mediated, GluN1/GluN2B/GluN2D-mediated, or GluN1/GluN2D-mediated excitatory postsynaptic currents in native neuronal systems (Figure 2A). The average decay constants for all cells were clustered closely together near the center of the decay range for each bin in both rats (Supplementary Figure 1A) and humans (Supplementary Figure 1B), supporting the use of these decay rate boundaries. Using this binning approach, we identified exponentially decaying mEPSC events that had a decay range between 1-20 ms (Figure 2B, yellow leftmost bars of graphs) with average decay constants that clustered around 10-15 ms for recordings from male and female, rat and human lamina I neurons (Supplementary Figure 1Ai,Bi; Supplementary Figure 2A,3A), which is consistent with AMPAR-dominated excitatory synaptic responses (Bourinet et al., 2014; Lin et al., 2002; S. J. Liu & Cull-Candy, 2002). The average decay constants for the mEPSCs in the 20-100 ms bin (Figure 2B, second from left yellow bars) were mainly distributed around 50-70 ms (Supplementary Figure 1Aii, Bii; Supplementary Figure 2B,3B), which is consistent with previously published average decay constants for GluN2A-mediated excitatory postsynaptic currents (Kumar & Huguenard, 2003). Similarly, mEPSC events with decay rates in the 100-200 ms and 200-400 ms bins (Figure 2B, middlemost two bins) and in the 400-1000 ms and 1000+ ms bins (Figure 2B, rightmost two bins) corresponded to GluN2B– and GluN2D-containing synaptic NMDAR responses, respectively (Supplementary Figures 1-3; for review, see (Paoletti et al., 2013)). Given accumulating evidence for triheterosynaptic NMDARs at native neuronal synapses, it should be noted that the 100-200 ms and 400-1000 ms bins likely also include GluN2A/GluN2B– and GluN2B/GluN2D triheteromeric receptor mediated synaptic responses, respectively (Gibb et al., 2018).

For both rat and human lamina I neurons, we found that individual neurons had a percentage of events that fell within the majority of the distinct AMPAR– and NMDAR-mediated mEPSC decay constant bins (n = 9 male rat neurons, n = 9 female rat neurons, n = 12 male human neurons, n = 4 female human neurons; Figure 2B, Supplementary Figure 1). A two-way ART ANOVA within species revealed no statistically significant differences in percent events per bin between sex (p = 1.0 for rat; p = 1.0 for human; Figure 2B); however, the percent frequency of mEPSC events in each of the decay constant-defined bins significantly differed across bin type (p = 1.5E-21 for rat; p = 7.8E-11 for human; Figure 2B). For a list of pairwise comparisons of percent frequency of mEPSC events in each of the decay constant-defined bins for both rats and humans, see Tables 1 and 2, respectively. Furthermore, in rats (sexes combined), the distribution of the percent frequency of the mEPSC decay constant-defined bins was approximately symmetrical (p = 0.45 ± 0.23, Figure 2B), while the event distribution (sexes combined) was moderately leftward skewed for human lamina I neurons (p = 0.92 ± 0.25, Figure 2B). This suggests that although both rat and human lamina I neurons exhibit wide heterogeneity in the decay constants of individual mEPSCs that is conserved across sex, the relative contribution of distinct NMDAR subunits in mediating this synaptic heterogeneity may differ between rats and humans.

### There is a decrease in the relative contribution of GluN2B-mediated NMDAR responses at human lamina I synapses compared to rats

Using GluN2 subunit-specific pharmacological antagonists, we have previously shown that GluN2B-containing NMDARs dominate mEPSCs in lamina I neurons from adult male rats (Hildebrand et al., 2014). Administration of the GluN2B-selective antagonist, Ro25-6981, blocked approximately half of synaptic NMDAR current charge transfer and the average decay constant of this Ro25-6981-sensitive NMDAR mEPSC component was 281 ms (Hildebrand et al., 2014). Consistent with this, here we found a higher proportion of putative GluN2B-dominated mEPSCs in the 200-400 ms decay constant bin compared to putative GluN2A-dominated mEPSCs in the 20-100 ms bin for recordings on lamina I neurons of both male and female rats, although this difference was not significant (Figure 2B, Table 1). In a striking divergence from our findings in rats, we found that mEPSCs from human lamina I neurons had a significantly (p = 1.50E-5, sexes combined) higher fraction of GluN2A-dominated NMDAR responses (20-100 ms mEPSC decay constant bin) compared to GluN2B-dominated NMDAR mEPSCs (200-400 ms decay constant bin) for both males and females (Figure 2B, Table 2).

To complement our decay constant binning analysis using GluN2 subunit-associated decay constant ranges (Figure 2A), we next used an unbiased Gaussian distribution approach to plot the probability densities of mEPSC decay constants for all cells in each group (male and female rats, male and female humans; Figure 2C). Similar to the decay constant binning approach, three distinct populations of mEPSC events were identified, with peak probability ranges at decay constants that generally corresponded to AMPAR/GluN2A-, GluN2B-, and GluN2D-dominated synaptic responses (Brickley et al., 2003; Hansen et al., 2014; Hildebrand et al., 2014; S. J. Liu & Cull-Candy, 2002; Paoletti et al., 2013). This distribution of three distinct excitatory synaptic decay rate populations was conserved across sex in both rats and humans (Figure 2C). As found with binning analysis (Figure 2B), Gaussian distribution analysis revealed that the ratio of faster (AMPAR/GluN2A-like) to intermediate (GluN2B-like) decaying excitatory synaptic events was higher in humans compared to rats (Figure 2C). To ensure that differences in the relative contribution of specific NMDAR subunits, as measured by decay constant, are not due to differential levels of space clamp between individual synapses and cells(Rall & Segev, 1985), we measured and compared the initial rise slope of average mEPSCs within each decay constant range for each recorded cell. We found no statistically significant difference between the fast component mEPSC rise slope by species or decay constant range (Supplementary Figure 4; p = 0.936, sexes combined).

To validate the above biophysical analysis approaches, we leveraged our previous GluN2-specific pharmacological strategies and findings from rat lamina I neurons (Hildebrand et al., 2014) for proof-of-concept pharmacological experiments in a human spinal cord slice. We initially recorded baseline mEPSCs at +60 mV from a female human lamina I neuron during perfusion of control ACSF. Following 13 minutes of control recording, we sequentially added 1 µM Ro25-6981 (Fischer et al., 1997) to block GluN2B-mediated NMDAR responses, 1 µM Ro25-6981 + 10 µM DQP-1105 (Acker et al., 2011) to block GluN2B and GluN2D, and 1 µM Ro25-6981 + 10 µM DQP-1105 + 10 µM TCN-201 (Hansen et al., 2012) to block GluN2B-, GluN2D-, and GluN2A-mediated NMDAR currents, with each new antagonist added after 20-30 minutes of recording (Figure 3A). We used our decay constant binning approach to identify the percentage of mEPSCs within each of the six decay constant bins for each treatment (Figure 3B).

**Figure 3.**
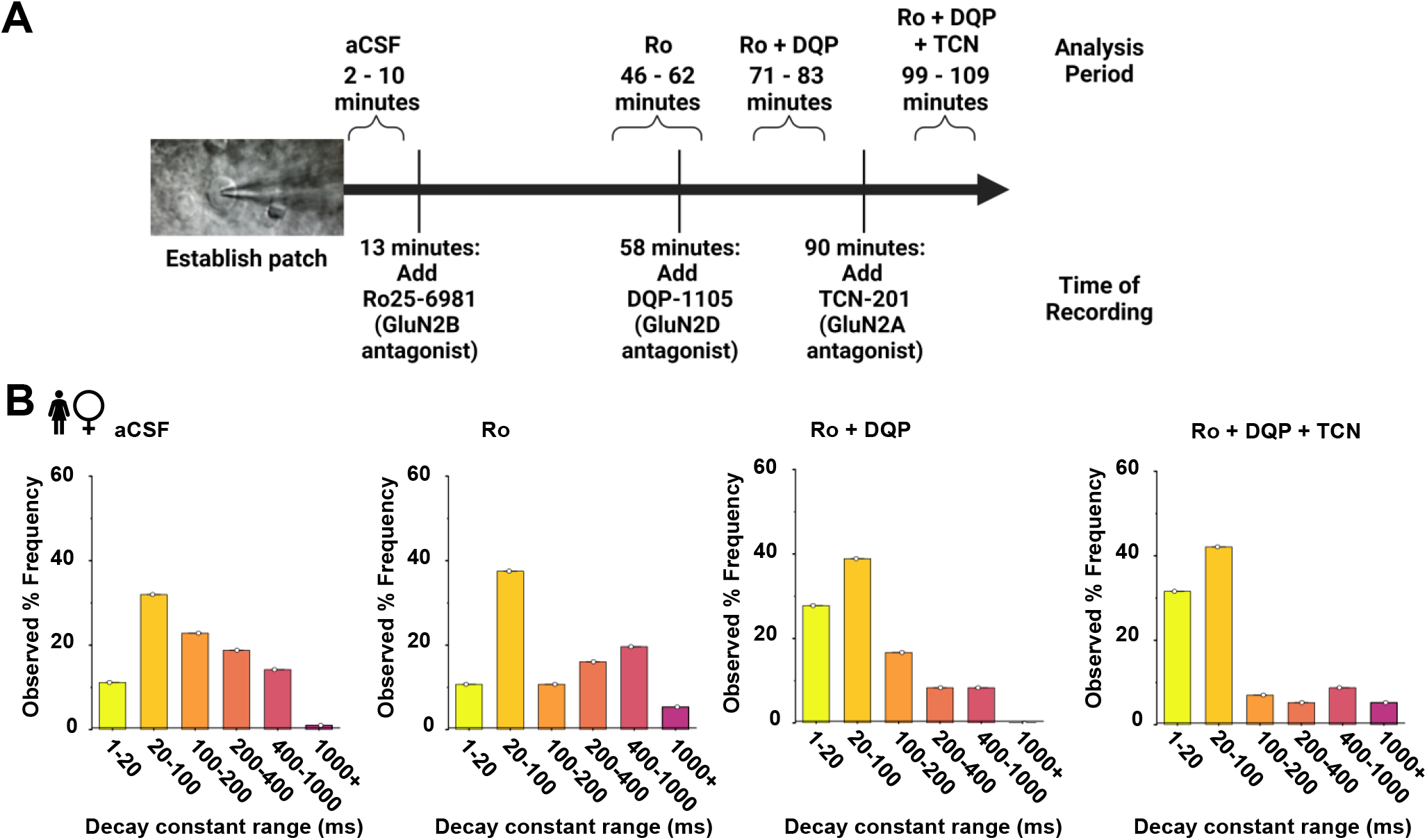
Administration of antagonists against specific GluN2 subunits reduces the proportion of mEPSCs within the corresponding decay constant ranges in a recording of a female human lamina I neuron. **A**) Diagram of pharmacological treatment timeline (below) and associated periods of mEPSC analysis (top). After establishing whole-cell voltage-clamp recording of an individual lamina I neuron, a control aCSF period was recorded, followed by subsequently adding 1 µM Ro25-6981, then 10 µM DQP-1105, and finally 10 µM TCN-201. **B)** Histograms of the proportion of mEPSCs in each decay constant bin during administration of control aCSF (left), 1 µM Ro25-6981 to block GluN2B NMDARs (middle left), 1 µM Ro25-6981 + 10 µM DQP-1105 to block GluN2B and GluN2D NMDARs (middle right), and 1 µM Ro25-6981 + 10 µM DQP-1105 + 10 µM TCN-201 to block GluN2B, GluN2D, and GluN2A NMDARs.

Under control conditions, the highest percent of events occurred in the GluN2A-like 20-100 ms bin (32.0%), followed by 22.8% of events in the GluN2A/GluN2B-like 100-200 ms bin and then and 18.8% of events in the GluN2B-like 200-400 ms bin (Figure 3B, left). With administration of the activity-dependent GluN2B antagonist Ro25-6981 (Fischer et al., 1997) for 33 minutes, the relative contribution of putative GluN2B-mediated synaptic NMDAR responses started to decrease (10.7% in 100-200 ms bin, 16.1% in 200-400 ms bin), while the relative proportion of putative GluN2A-like event increased (37.5% in 20-100 ms bin) (Figure 3B, middle left).

Following this, continued perfusion of Ro25-6981 with co-treatment of the GluN2D-antagonist DQP-1105 (Acker et al., 2011) for 13 minutes induced a continued decline in putative GluN2B events in the 200-400 ms bin (8.33%) (Figure 3B middle right). The addition of DQP-1105 during this period reduced putative GluN2B/GluN2D– and GluN2D-mediated synaptic responses in the 400-1000 ms and 1000+ ms decay constant bins, respectively, from 19.6% and 5.4% during initial perfusion of Ro25-6981 only (Figure 3B middle left) to 8.3% and 0% during perfusion of Ro25-6981 and DQP-1105 (Figure 3B middle right). Finally, addition of the GluN2A antagonist TCN-201 (Hansen et al., 2012) for 9 minutes during the continued perfusion of Ro-25-6981 and DQP-1105 induced a decline in putative GluN2A/GluN2B-mediated synaptic NMDAR responses in the 100-200 ms bin from 16.7% to 7.0%, with no decrease in the proportion of events in the 20-100 ms bin during this initial 10 min administration of TCN-201 (Figure 3B right). Complete blockade of GluN2A NMDARs by continued application of TCN-201 was not able to be reached, as this robust human lamina I neuron was lost after 115 minutes of patch-clamp recording. These current and previous (Hildebrand et al., 2014) pharmacological experiments validate that the designated decay constant bins accurately correspond to the assigned contributing AMPAR and GluN2 subunits shown in Figure 2A for recordings on rat and human lamina I neurons. Thus, from the above biophysical and pharmacological experiments, we conclude that the relative contribution of GluN2B versus GluN2A to spontaneous synaptic NMDAR responses in lamina I neurons decreases in humans compared to rats.

### mEPSC amplitude is conserved across sex, with more variability and larger slow-decaying GluN2D-like events in humans compared to rats

Our initial rodent-optimized mEPSC analysis criteria (Hildebrand et al., 2014) excluded numerous large amplitude spontaneous synaptic events that were observed in preliminary recordings from male human lamina I neurons (Dedek et al., 2019). Here, we used an unbiased analysis approach with no prescribed mEPSC upper amplitude limit to investigate if the amplitudes of distinct subtypes of mEPSC events are conserved between sexes in rat and human lamina I neurons. In examining the peak mEPSC amplitude averaged within each cell across decay constant-defined bins, we found a tight distribution of amplitudes in male and female rat mEPSC responses (Figure 4A; Supplementary Figure 5A). In male rats, the average peak amplitude within each bin ranged from 13.9±1.7 to 25.5±6.0 pA, while in female rats the range was from 13.1±0.6 to 20.5±3.4 pA (Figure 4A). A two-way ART ANOVA revealed no statistically significant differences in amplitude between sex or decay constant range in rats (p = 0.90 for comparison by sex, and p = 0.16 for comparison by decay constant-defined bin; male rat n = 9 neurons, female rat n = 9 neurons; Figure 4A). We observed higher variability in individual and averaged human mEPSC peak amplitudes, which ranged from 16.8±4.2 to 121±54 pA in males and 11.3±2.7 to 52.0±17.6 pA in females (Figure 4B; Supplementary Figure 5B). A two-way ART ANOVA within species revealed no statistically significant differences in amplitude between sex or decay constant range (p = 0.72 for comparison by sex, and p = 0.32 for comparison by decay-constant defined bin; male human n = 12 neurons, female human n = 4 neurons; Figure 4B). Although direct statistical comparisons cannot be made due to differences in the preparation, the average peak amplitudes of NMDAR mEPSCs in human lamina I neurons were sometimes substantially greater than that found in rats. This increased variability was more pronounced for the higher-powered male lamina I neuron dataset (Figure 4B). We also found that the slowest over-1000 ms decay constant-defined bin corresponding to GluN2D-mediated synaptic NMDAR responses had visibly higher average amplitudes compared to rats in both sexes (Figure 4B).

**Figure 4.**
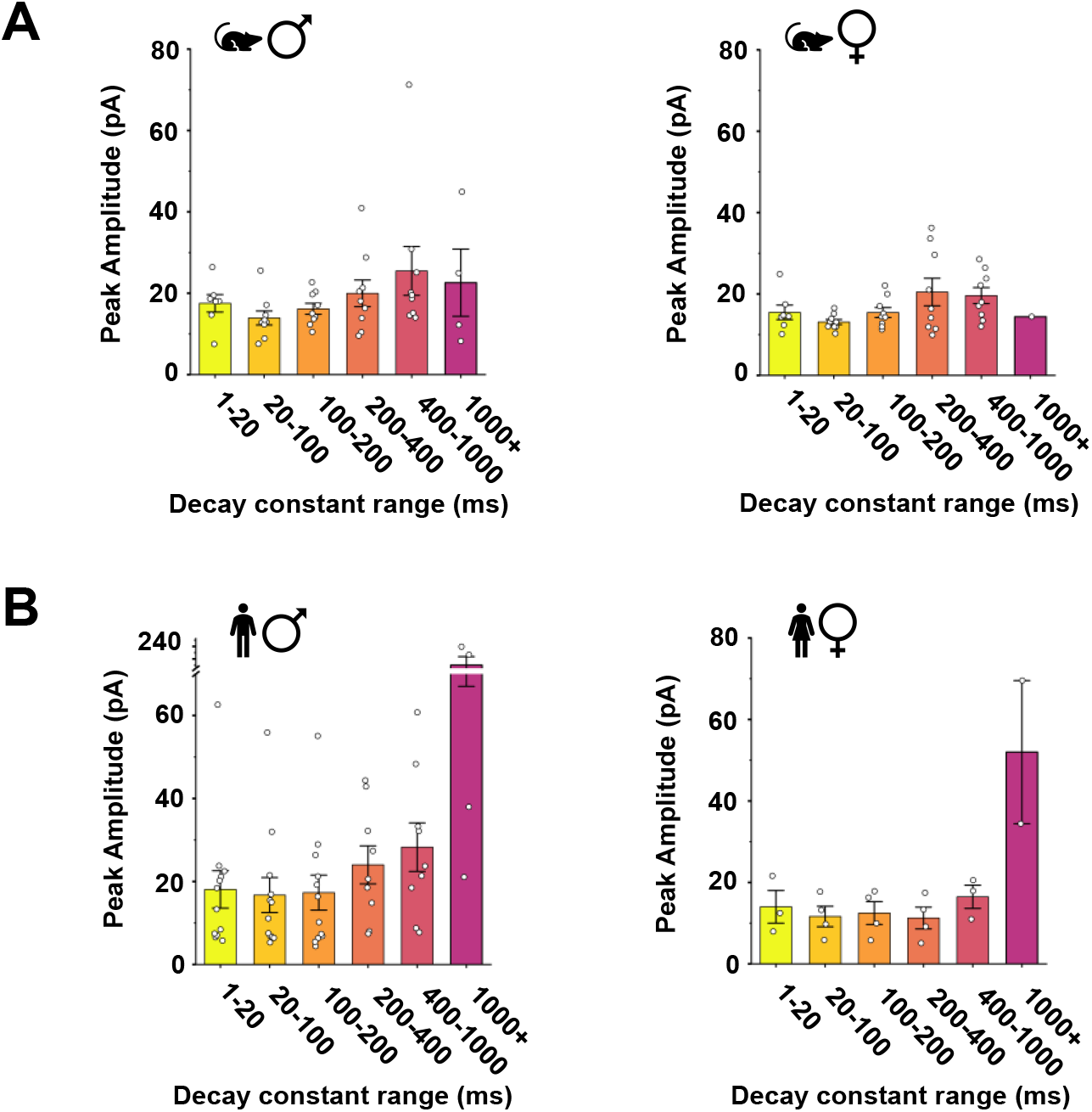
mEPSC amplitude is conserved across sex within species, with larger amplitudes of GluN2D-like events observed in humans compared to rats. **A,B**) Plots of average peak amplitude, in pA, by cell for all decay constant range bins in male (left) and female (right) rats (**A**) and humans (**B**). Male rat n = 9, female rat n= 9, male human n = 12, female human n = 4.

## Discussion

Given the urgent need to understand the physiological properties of pain processing circuits across sex as well as from rodents to humans, we used patch clamp recordings to systematically characterize the biophysical properties of excitatory synaptic currents in viable rat and human lamina I neurons. We found that averaging synaptic mEPSCs masked a surprising degree of intrinsic variability within individual lamina I neurons, and so developed an unbiased analysis approach that measured the decay constant and peak amplitude for each single synapse mEPSC event. Using this approach, we identified a wide range of decay constants for mEPSCs within individual lamina I neurons, which corresponded to distinct AMPAR– and GluN2-NMDAR subunit-mediated synaptic responses. This intraneuronal heterogeneity in mEPSC properties was conserved across sex for both rat and human lamina I neurons. Importantly, a combination of biophysical and pharmacological experiments led to the discovery that GluN2B-mediated NMDAR synaptic responses are most prevalent in male and female rats, while GluN2A-mediated NMDAR responses dominate in male and female human lamina I neurons. Although no significant differences in the amplitudes of mEPSCs were found across sex for both rats and humans, we found higher variability in mEPSC amplitudes in human lamina I neurons, with visibly higher-amplitude GluN2D-mediated mEPSC events in humans compared to rats.

Stimulation of distinct neuroanatomically-defined inputs (Banerjee et al., 2014; Ito et al., 2000; Kumar & Huguenard, 2003; Larsen & Sjöström, 2015; Otmakhova et al., 2002) has been shown to activate different NMDAR-dependent synaptic population responses and drive synapse-specific mechanisms of plasticity within cortical and hippocampal neurons. Here, we find an even larger degree of heterogeneity in AMPAR– and NMDAR-driven synaptic currents within individual rat and human lamina I neurons, through analysis of mEPSCs that represent the sum of all possible quantal postsynaptic responses within a neuron (Hildebrand et al., 2014). This synaptic heterogeneity may correspond to the divergent array of inputs that lamina I neurons receive from distinct sensory neuron afferents, local excitatory and inhibitory spinal interneurons, and descending efferents. Consistent with our findings, presynaptic afferent-evoked excitatory responses in male rats also exhibit considerable heterogeneity that corresponds to different constituent NMDAR subunits (Hildebrand et al., 2014; Pitcher et al., 2023; Tong & MacDermott, 2014). In tour de force recordings of minimally-evoked EPSCs in male rat lamina I neurons, Pitcher and colleagues combined unbiased principal component and hierarchical analysis with GluN2 subunit-specific antagonists to demonstrate that afferent-evoked *unitary* EPSCs have either fast (52±3 ms), intermediate (315±11 ms), or slow (1219±69 ms) decay kinetics, which correspond to GluN2A, GluN2B, or GluN2D NMDARs, respectively (Pitcher et al., 2023). Importantly, activation of these distinct decaying quantal postsynaptic responses drives different degrees of temporal summation and is correlated with divergent lamina I neuron firing patterns (Pitcher et al., 2023). Our findings here show that this newly discovered heterogeneity in NMDAR function at single synapse resolution is conserved across sex for both rats and humans.

Despite synapse-specific heterogeneity in lamina I NMDARs being found in both rats and humans, the relative contribution of underlying GluN2 subunits diverges across species. Both findings from mEPSC analysis here and recordings of afferent-evoked unitary EPSCs demonstrate (Pitcher et al., 2023) that intermediate-decaying, GluN2B-mediated NMDAR responses are most prevalent at rat lamina I synapses. In contrast, we find that fast-decaying GluN2A-mediated synaptic responses dominate in male and female human lamina I neurons. However, it should be noted that there were differences in the rat versus human patch-clamp recording preparations, such as the slices being from lumbar spinal cord in the parasagittal plane for rodents while being primarily from sacral spinal cord tissue in the transverse plane for humans. Nonetheless, this species difference has important potential clinical implications. In many rodent models of chronic pain, GluN2B antagonists have been shown to reverse behavioural hypersensitivity (Bourinet et al., 2014), while these promising GluN2B-targetting preclinical candidates have generally failed in human clinical trials (W. Liu et al., 2020). One reason for this lack of translation could potentially be due to a less pronounced role for GluN2B NMDARs in human dorsal horn pain processing.

Recent experimental evidence has shone a light on the GluN2D NMDAR subunit as a critical mediator of synaptic signalling in spinal pain processing neurons. Although GluN2D expression is limited in the mature brain (Paoletti et al., 2013), this unique NMDAR subunit is highly expressed and localized to the superficial dorsal horn of adult male and female rats and humans (Armstrong et al., 2022; Temi et al., 2021). GluN2D-containing NMDARs have slower deactivation kinetics, less sensitivity to blockade by external Mg^2+^ ions, and higher affinity for glutamate agonist binding, all enabling enhanced temporal and spatial summation (Paoletti et al., 2013). Here we show for first time that prolonged GluN2D-dominated synaptic NMDAR events are found across male and female rat and human lamina I neurons, with a potential for larger GluN2D-mediated NMDAR responses at human lamina I synapses. As slow decaying GluN2D-mediated synaptic responses have been shown to increase integrative capacity in lamina I neurons of adult male rats (Pitcher et al., 2023), it will be critical to investigate the impacts of this relatively spinal-specific NMDAR subtype on dorsal horn plasticity and network processing.

In preclinical investigations into the roles of specific molecular determinants (such as NMDAR subtypes) in pain processing, there is a large gap between target functions and associated pain behaviors. Understanding how these targets influence local and global signalling in spinal pain circuits remains poorly understood but is critical for potential translation given the degeneracy found within these networks (Ratté & Prescott, 2016). For example, what is the functional impact of the intraneuronal synaptic heterogeneity found here on differential mechanisms of plasticity and excitability in defined dorsal horn nociceptive networks? With the development of novel human tissue physiological assays (Dedek et al., 2019, 2022), we can now test whether spinal circuit-level mechanisms of pain processing are conserved or diverge across sex and species and investigate how manipulating specific targets with novel therapeutic candidates alters excitability in human spinal nociceptive networks.

## Methods

### Animals

All animal experiments followed the guidelines and policies provided by the Canadian Council for Animal Care, Carleton University, and the University of Ottawa Heart Institute. Both male and female adult (3-4 months old) Sprague-Dawley rats were used. All rats were provided by Charles River Laboratories. The rats were housed in same sex-pairs with a 12-hour day-night cycle and had access to water and food *ad libitum*.

### Rat Spinal Cord Isolation

Rats were deeply anesthetized by intraperitoneal injection of 3g/kg urethane (Sigma), and vertebral corpectomy was used to rapidly dissect spinal cords. The isolated spinal cords were directly transferred into ice-cold oxygenated protective sucrose solution containing: 92 mM NaCl, 50 mM sucrose, 26 mM NaHCO_3_, 15 mM D-glucose, 7 mM MgSO_4_, 5 mM KCl, 1.25 mM NaH_2_PO_4_, 1 mM kynurenic acid, 0.5 mM CaCl_2_, and was bubbled with 95% O_2_/5% CO_2_. The L3-L6 lumbar region was viewed under a dissection microscope, and dorsal and ventral roots were removed with forceps and vannas spring scissors. Parasagittal slices of 300 µm thickness, mainly including the lateral half of each dorsal horn, were cut with a Leica VT1200S vibratome at 2.75 mm amplitude and 0.1-0.3 mm/s blade advance speed. After slicing, slices were incubated in 34°C kynurenic acid-free protective solution for 40 min to promote recovery and to wash out kynurenic acid. Previous control experiments have shown no difference in NMDAR-mediated synaptic responses in lamina I neurons from slices that were sectioned in protective solution with or without kynurenic acid, provided that slices had recovered in kynurenic acid-free protective solution (data not shown). Following kynurenic acid washout and recovery, the incubation chamber was removed from the heated water bath and allowed to passively cool to room temperature before proceeding with experiments.

### Human Donors

Human organ donor studies were approved by the Ottawa Health Science Network Research Ethics Board. Neurologic and cardiac determination of death donors were identified through the Trillium Gift of Life Network. Human spinal tissue was collected from male and female human donors between the ages of 18-70 years old. Candidates for donation were screened for communicable diseases (such as HIV/AIDS and syphilis) and conditions that could negatively affect the health of organs, such as morbid obesity. Donors with known medical conditions that would compromise the health of the central nervous system were eliminated from the study. The primary cause of death was an interruption of blood flow to the brain by ischemia or hemorrhage.

### Human Spinal Cord Isolation

Hypothermia was induced by placing human donors on a cooling bed. Organs were removed for donation and then spinal cord isolation by vertebral corpectomy was performed within 3 hours after aortic cross-clamp or organ flushing. The lumbar, and/or sacral regions of the spinal cord were isolated in the operating room and placed into ice-cold oxygenated protective solution (see above for composition) solution containing 1 mM kynurenic acid. Careful removal of all meninges was performed under a dissection microscope. Transverse spinal 500 µm thick slices were cut with a Leica VT1200S vibratome at 2.75 mm amplitude and 0.1 mm/s speed during sectioning through the dorsal horn. Similar to rat slices, the kynurenic acid-free protective solution was used to wash out kynurenic acid for a 40-minute incubation at 34 °C, followed by a passive return to room temperature.

### Electrophysiology Recordings

Spinal cord slices were viewed under brightfield optics. Lamina I neurons were identified as being situated dorsal to the substantia gelatinosa and within 50 µm of the overlying superficial white matter tracts. In some human spinal cord slices, the outer edge of the substantia gelatinosa, visualized under low magnification, was used to orient when no white-matter tracts were present. As previously described (Dedek et al., 2019, 2022; Hildebrand et al., 2016), the extracellular recording solution contained 125 mM NaCl, 26 mMNaHCO_3_, 20 mM D-glucose, 3 mM KCl, 2 CaCl_2_, 1.25 mM NaH_2_PO_4_, and 1 MgCl_2_, as well as the blockers for voltage-gated Na^+^ channels (500 nM TTX), voltage-gated Ca^2+^ channels (10 μM Cd^2+^), glycinergic currents (10 μM strychnine), and GABAergic currents (10 μM bicuculline). We fire-polished borosilicate glass patch-clamp pipettes with 6-12 MΩ resistance (Sutter Instruments). The internal patch solution had a pH of 7.25, osmolarity of approximately 295 mOsm, and contained 105 mM Cs-gluconate, 17.5 mM CsCl, 10 mM BAPTA or EGTA, 10 mM HEPES, 2 mM MgATP, and 0.5 mM Na_2_GTP. Whole-cell patch was established at –60 mV. We only included neurons that showed access resistance below 35 MΩ and leakage current above –100 pA. We slowly increased the holding potential to +60 mV to remove the Mg^2+^ block from NMDARs. Outward miniature excitatory postsynaptic currents (mEPSCs) were measured by Clampfit 11.1.0.23 (Molecular Devices).

### Pharmacology

Pharmacology experiments were performed by acutely perfusing selective NMDAR antagonists onto a slice, following the control ACSF period. (αR,βS)-α-(4-Hydroxyphenyl)-β-methyl-4-(phenylmethyl)-1-piperidinepropanol maleate (Ro25-6981) (Fischer et al., 1997), 4-(5-(4-bromophenyl)-3-(6-methyl-2-oxo-4-phenyl-1,2-dihydroquinolin-3-yl)-4,5-dihydro-1H-pyrazol-1-yl)-4-oxobutanoic acid (DQP-1105) (Acker et al., 2011), and (3-chloro-4-fluoro-N-[(4-([2-(phenylcarbonyl)hydrazino]carbonyl) phenyl)methyl]benzene-sulfonamide) (TCN-201) (Hansen et al., 2012)were obtained from Tocris Bioscience. Stock solutions were prepared in DMSO. Antagonists were sequentially added to ACSF to block GluN2B (1 µM Ro25–6981; (Fischer et al., 1997)), GluN2B and GluN2D (1 µM Ro25–6981 + 10 µM DQP; (Acker et al., 2011)), and finally GluN2B, GluN2D and GluN2A (1 µM Ro25–6981 + 10 µM DQP + 10 µM TCN-201; (Hansen et al., 2012)). Antagonists were perfused onto the slice for 33 minutes for Ro25-6981, 13 minutes for DQP, and 9 minutes for TCN-201 before analysis for each respective analysis period.

### GluN2B-Focussed Detection and Analysis

The threshold search feature within Clampfit 11.2 (Molecular Devices) was used to detect positive-going mEPSC events. The first two minutes of recording at +60 mV were omitted to allow the neuron to acclimate. The following eight minutes of the recording were analyzed to ensure consistency across recordings and to prevent confounds due to potential run-up or run-down due to potential changes in the intracellular milieu during longer term recordings. Pre– and post-trigger lengths were set to 250 ms and 1 s, respectively. Events with durations less than 40 ms and amplitudes less than 10 pA were considered noise. Selection criteria for mEPSCs included: no events that overlap, no events that completely decay within 100 ms, no outlier events with an amplitude >200 pA, and events must decay to at least 50% of their overall amplitude by 500 ms. mEPSC traces were averaged together by sex and species in Clampfit 11.2.

### Unbiased Event Detection and Analysis

The threshold search feature within Clampfit 11.2 was used to detect positive-going mEPSC events. The first two minutes of recording at +60 mV were omitted to allow the neuron to acclimate. The following eight minutes of the recording were analyzed to ensure consistency across recordings and to prevent confounds due to potential run-up or run-down due to potential changes in the intracellular milieu during longer term recordings. Pre– and post-trigger lengths were set to 500 ms and 2 s, respectively. Events with durations less than 10 ms and amplitudes less than twice the peak-to-peak background signal were considered noise. To prevent any bias for larger and more clear events, all events detected by the search feature with any amplitude and any exponential decay times were accepted as long as the rise time was less than the decay time, discrete events did not overlap with each other, and there was no major baseline shift. Visual inspection was done during event detection to eliminate events where the baseline shift was greater than the amplitude, with more specific elimination done in the later fitting step.

All selected events were aligned in Clampfit 11.2. Exponential one-term standard fitting with the Chebyshev search method and default seed values were applied for each event with the formula: *f*(*t*) = *A* ∗ *e*^−*t*/τ^ + C. Cursor 1 and cursor 2 were manually aligned to define the fitting range with the following approach: excitatory postsynaptic currents can have both fast AMPAR-like and slow NMDAR-like components, with corresponding fast and slow decay times, respectively (Lester et al., 1990; Sah et al., 1990). Cursor 1 was placed so that only AMPAR or NMDAR-dominated mEPSCs were analyzed for each event. In the case of the mEPSC showing the properties of an AMPAR-only current with a fast rise and a steep decay that returned to baseline within 40 ms, cursor 1 was placed just after the peak, approximately 0.5-2 ms from event onset. If the mEPSC displayed properties of NMDARs only, with relatively slow increase and decay time, cursor 1 was placed approximately 5-10 ms past the decay inflection, typically approximately 20-200 ms from event onset. If the event displayed properties of both AMPAR and NMDAR currents, cursor 1 was placed after the steep decline of AMPAR current to focus on the slow NMDAR component’s decay rate only. Cursor 2 was set to a time point where the decay event was either plateaued at or around the baseline level. Throughout the analysis, “amplitude” refers to the “A” value obtained by the fitting, rather than the amplitude of the whole event, which can be distorted by noise or may differ from the fitted event due to the AMPAR-mediated component. If the fitted line deviated more than 20% from the trace (determined by standard error / τ_decay_), the trace was excluded from analysis. Events with C/A greater than 30% were excluded (values obtained in the output of each fit). Events that gave non-physiological results such as negative amplitude and τ_decay_ values were also discarded. mEPSCs from each cell were separated into six bins based on τ_decay_. Bins corresponded to previously published ranges of decay constants for AMPAR-mediated, GluN1/GluN2A-mediated, GluN1/GluN2A/GluN2B-mediated, GluN1/GluN2B-mediated, GluN1/GluN2B/GluN2D-mediated, or GluN1/GluN2D-mediated excitatory postsynaptic currents in native neuronal systems (Figure 2A). When calculating the proportion of mEPSCs per cell in each bin, the number of mEPSCs was normalized to the overall number of events recorded in that cell.

### Rise Slope Analysis

For each cell, averaged mEPSCs were generated using all mEPSCs that fell within each decay constant range. Rise slope was measured by first determining the shortest rise time of all averaged mEPSCs (1.2 ms), then placing cursor 1 at the start of the steepest rise of the mEPSC, and cursor 2 1.2 ms from cursor 1. Clampfit was then used to measure the rise slope between cursors 1 and 2. This step was repeated for all averages of mEPSCs within each decay constant range for all cells.

### Statistical Analysis and Graphing

Data were analyzed using SPSS (IBM SPSS Statistics 27). For all tests, significance was set to α = 0.05. Normality tests for each variable were evaluated by the Kolmogorov-Smirnov test. Homogeneity of variance was tested with the Levene Statistic. When assumptions were satisfied, a two-factor ANOVA was used. When data violated assumptions of normality and homogeneity of variance, two-factor Aligned Rank Transform (ART) ANOVA was used in place of a two-factor ANOVA (Wobbrock et al., 2011a). The data were transformed using aligned rank transformation with ARTool (Wobbrock et al., 2011b, 2021). The transformed data were subsequently analysed using a one-factor ANOVA with Bonferroni post-hoc test in SPSS for each factor and possible interaction in the analysis (Elkin et al., 2021). Descriptive statistics were used to measure skewedness. Graphing was performed using Origin Pro (Northampton). All numbers reported with ± represent standard error of the mean (S.E.M.).

## Acknowledgements

We are grateful to organ donors and their families for their extremely generous gift; this study would not have been possible without them. We thank the intensive care unit and operating room staff at the Ottawa Hospital, Civic Campus for donating their time and energy to make the human tissue collection in this study possible. Thank you to the research staff and coordinators Suzan Chen, Lei Zhou, Angela Auriat, Jessica Parnell, Harleen Kaur, and Maitreya Patel. We are grateful to Graham M. Pitcher for his helpful feedback on this manuscript. A.D. received a Mitacs Accelerate Fellowship. This work was supported by an NSERC Discovery Grant (M.E.H.).

## Author contributions

A.D., C.D., and M.E.H. conceived of the study and designed experiments. A.D. and E.C.T. collected tissue. A.D., C.D., and M.E.H. performed electrophysiological experiments. A.D., E. T., and C.D. performed data analysis. A.D. and M.E.H. drafted the manuscript, with editing from E.T., C.D., J.S.M, J.L.K., and E.C.T.

## Data Availability

The data that support the findings of this study are available from the corresponding author, upon reasonable request.

## Figures

**Supplementary Figure 1.**
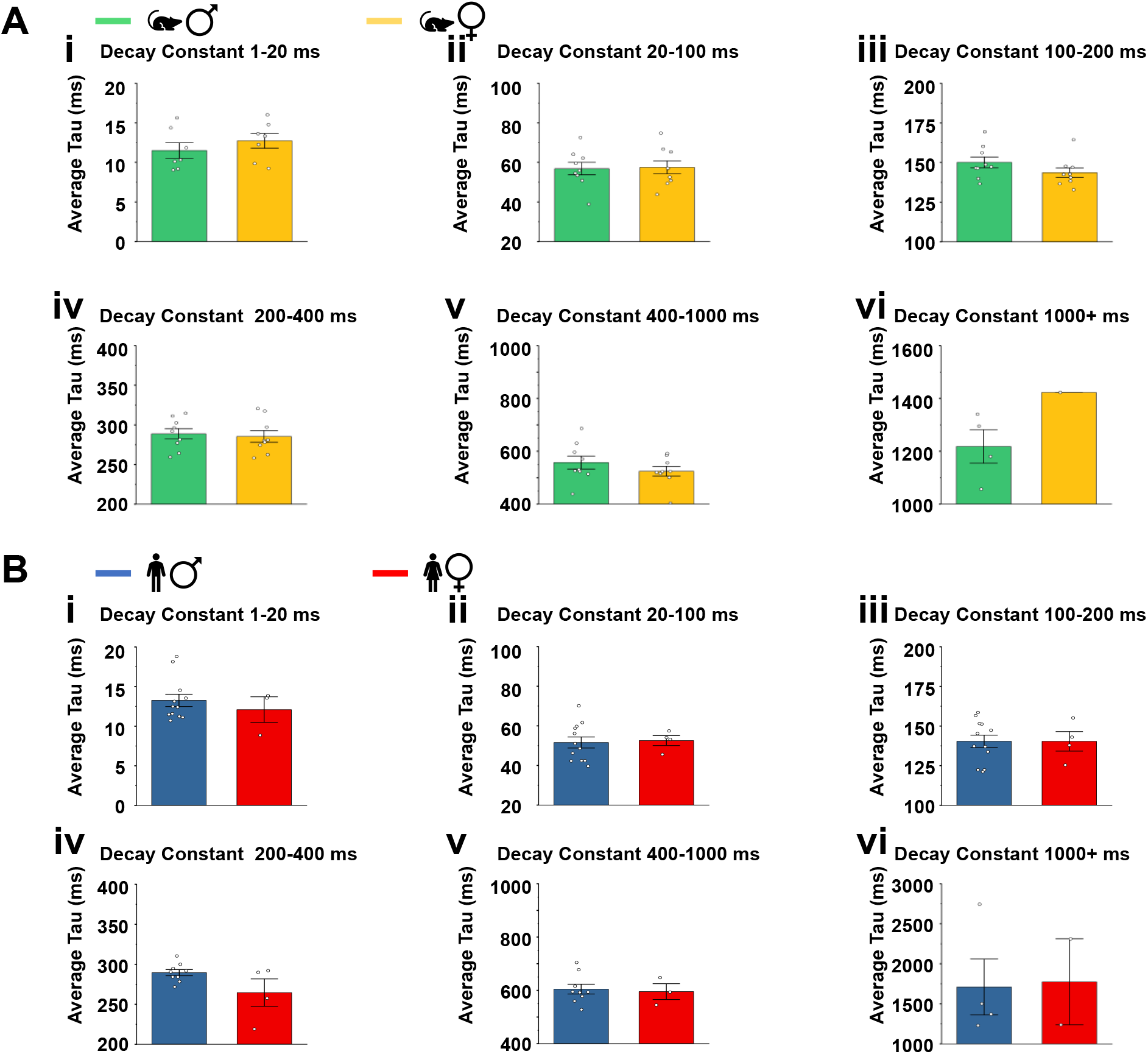
Average decay constants are clustered near the center of the defined decay constant ranges. **A**) The distribution of the average decay constants (tau) within each decay constant bin for cells recorded from male (green) and female (yellow) rat lamina I neurons. **B)** The distribution of the average decay constants (tau) within each decay constant bin for cells recorded from male (blue) and female (red) human lamina I neurons.

**Supplementary Figure 2.**
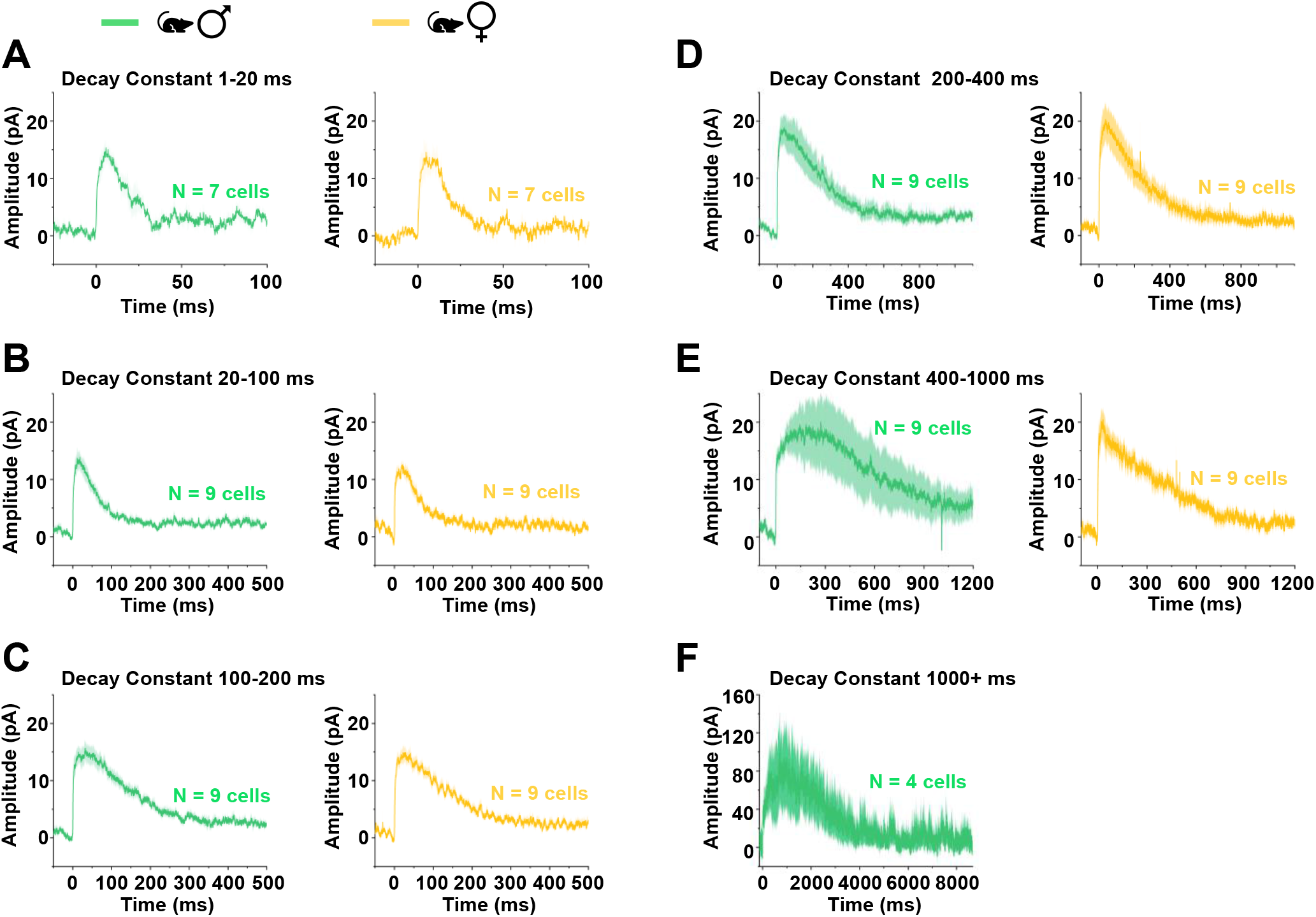
Rat lamina I neuron mEPSC traces averaged by decay constant range. **A-F**) Male (green) and female (yellow) rat lamina I neuron mEPSCs averaged for each decay constant range for all cells in the study, displayed ± SEM. N values are shown for each averaged plot.

**Supplementary Figure 3.**
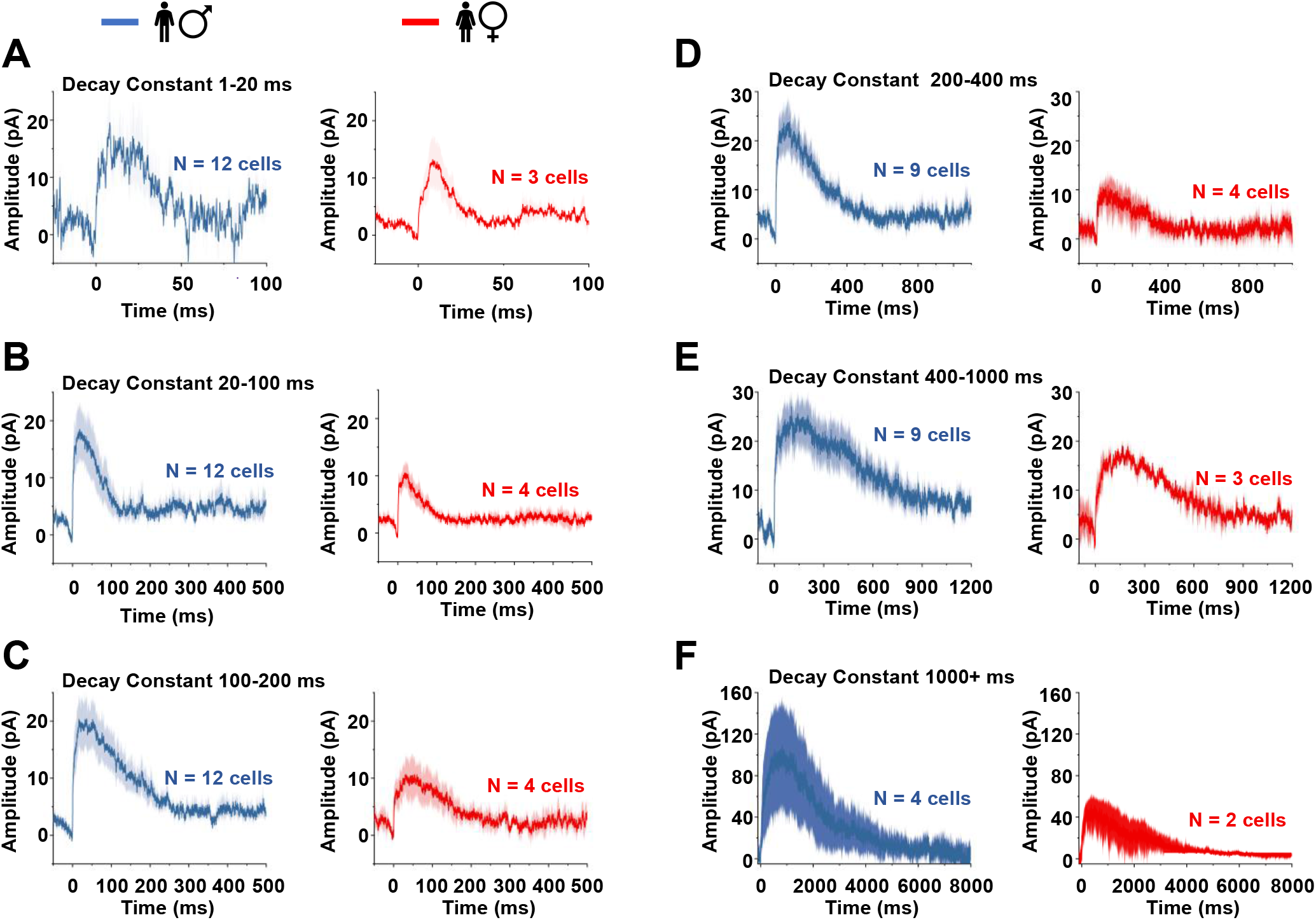
Human lamina I neuron mEPSC traces averaged by decay constant range. **A-F**) Male (blue) and female (red) human lamina I neuron mEPSCs averaged for each decay constant range for all cells in the study, displayed ± SEM. N values are shown for each averaged plot.

**Supplementary Figure 4.**
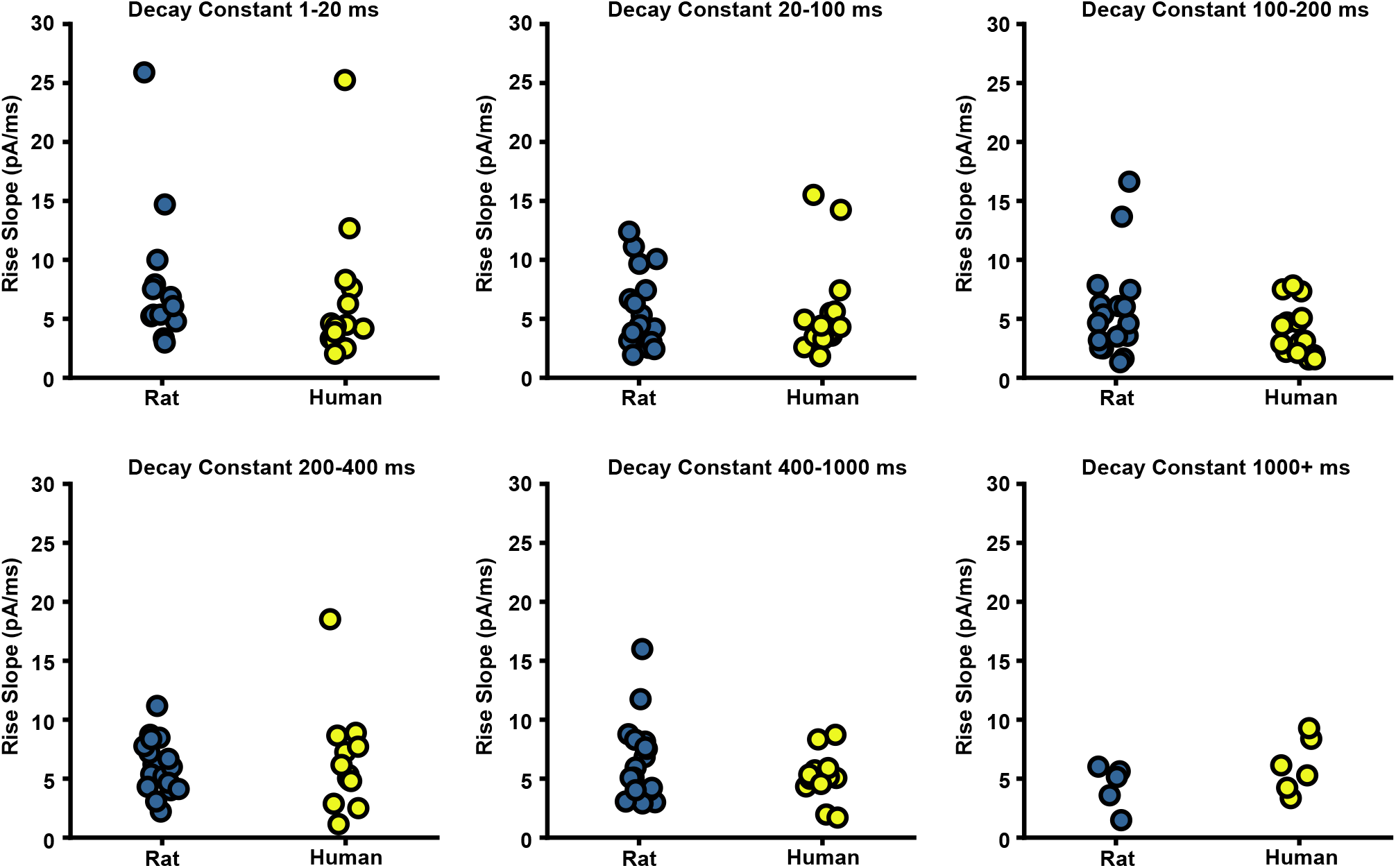
mEPSC initial rise slope does not differ by decay constant range or by species. Averaged mEPSC traces were generated for each decay constant range within each cell and were analyzed with sexes combined. 2-way ANOVA found no significant difference between initial mEPSC rise slope by species or by decay constant range (p = 0.936).

**Supplementary Figure 5.**
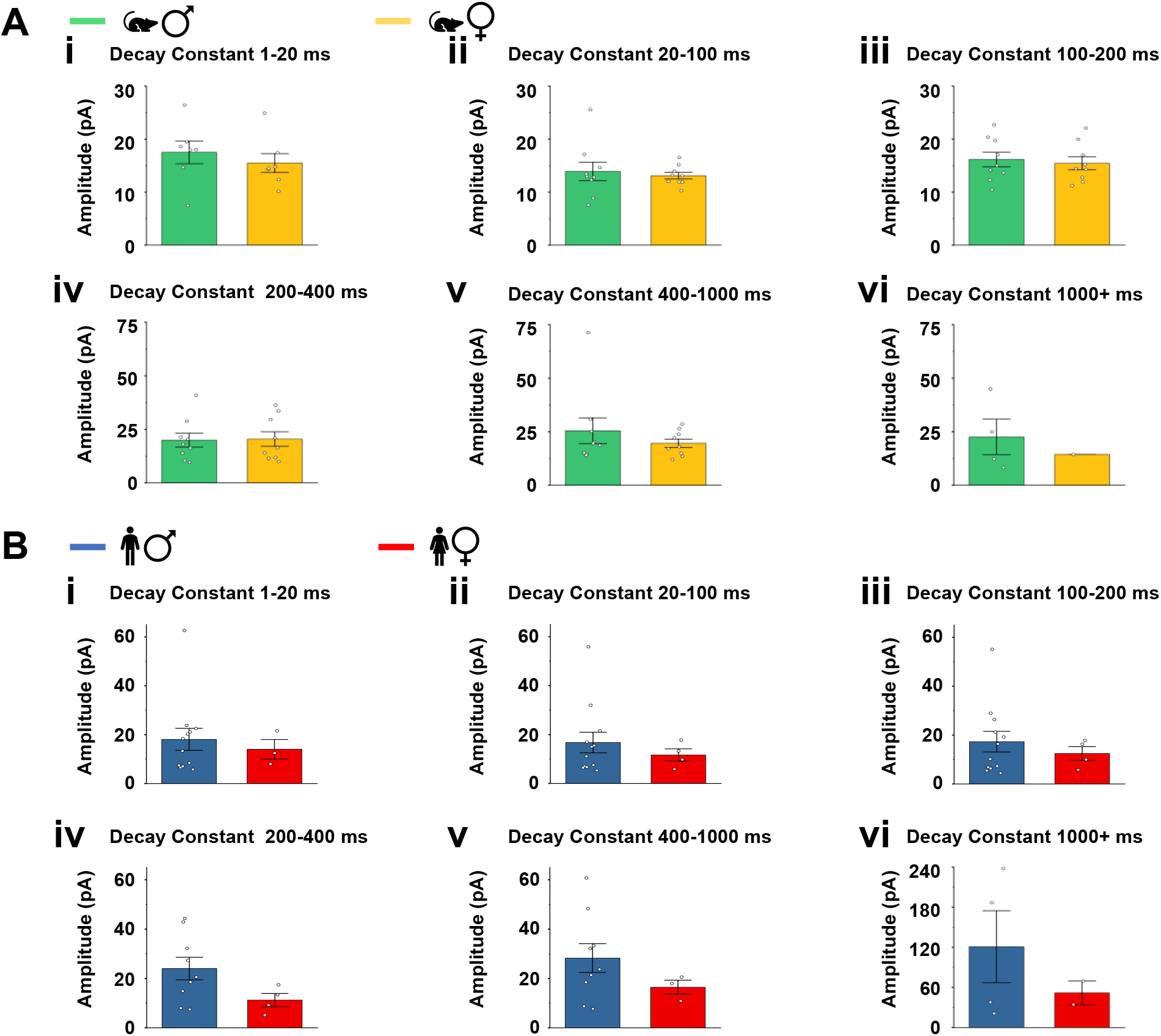
Lamina I mEPSC amplitudes are variable within decay constant-defined bins in rats and humans. **A,B**) Distribution of average amplitudes of all lamina I mEPSC amplitudes for rat **(A)** and human **(B)** cells within decay constant-defined bins (i – vi).

